# Characterization of OTUB1 activation and inhibition by different E2 enzymes

**DOI:** 10.1101/847806

**Authors:** Lauren T. Que, Marie E. Morrow, Cynthia Wolberger

## Abstract

OTUB1 is a highly expressed cysteine protease that specifically cleaves K48-linked polyubiquitin chains. This unique deubiquitinating enzyme (DUB) can bind to a subset of E2 ubiquitin conjugating enzymes, forming complexes in which the two enzymes can regulate one another’s activity. OTUB1 can non-catalytically suppress the ubiquitin conjugating activity of its E2 partners by sequestering the charged E2~Ub thioester and preventing ubiquitin transfer. The same E2 enzymes, when uncharged, can stimulate the DUB activity of OTUB1 *in vitro*, although the importance of OTUB1 stimulation *in vivo* remains unclear. In order to assess the potential balance between these activities that might occur in cells, we characterized the kinetics and thermodynamics governing the formation and activity of OTUB1:E2 complexes. We show that both stimulation of OTUB1 by E2 enzymes and noncatalytic inhibition of E2 enzymes by OTUB1 occur at physiologically relevant concentrations of both partners. Whereas E2 partners differ in their ability to stimulate OTUB1 activity, we find that this variability is not correlated with the affinity of each E2 for OTUB1. In addition to UBE2N and the UBE2D isoforms, we find that OTUB1 inhibits polyubiquitination activity of all three UBE2E enzymes, UBE2E1, UBE2E2, and UBE2E3. Interestingly, although OTUB1 also inhibits the autoubiquitination activity of UBE2E1 and UBE2E2, it is unable to suppress autoubiquitination by UBE2E3.

## Introduction

Ubiquitination is an essential post-translational modification that serves as a signal for a multitude of biological processes in eukaryotes. Ubiquitin is best known for its role in targeting substrates for proteasomal degradation *^1^*, but it also plays a role in nondegradative processes such as transcription regulation *^2^*, DNA damage repair signaling *^3, 4^*, and endosomal sorting *^5^*. The 76-amino acid ubiquitin protein is conjugated to a target substrate or to another ubiquitin molecule through the E1-E2-E3 enzyme cascade *^6^*. The C-terminus of ubiquitin (Ub) is conjugated to the E1 active site cysteine in an ATP-dependent manner and is then transferred to the catalytic cysteine of an E2 ubiquitin conjugating enzyme, forming an E2~Ub thioester intermediate, also referred to as a charged E2. Once conjugated to the E2, an E3 ligase catalyzes the transfer of ubiquitin to a substrate lysine, resulting in an isopeptide linkage between the ubiquitin C-terminus and the lysine ɛ-amino group. There are several classes of E3 ligases that differ in both structure and mechanism: the RING, HECT, RBR-like *^7^*, and the most recently discovered RCR enzymes *^8, 9^*. A majority of E3 ligases contain a RING finger domain, which serves as a mediator to bring the charged E2 and target substrate in close proximity to one another, promoting direct transfer of the ubiquitin to the substrate lysine *^9^*. Substrate proteins can be modified with a single ubiquitin (monoubiquitination), multiple single ubiquitin molecules, or a polyubiquitin chain in which ubiquitins are covalently linked to one another. Ubiquitin contains seven lysines, K6, K11, K27, K29, K33, K48 and K63, and an N-terminal amine (M1) to which another ubiquitin can be conjugated, resulting in eight different types of homotypic polyubiquitin chains *^10^*. Different chain types serve distinct biological roles: for example, K48-linked and K11-linked polyubiquitin chains target substrates for proteasomal degradation *^11^*, whereas K63-linked chains play non-degradative roles in DNA damage repair *^4^* and inflammatory signaling pathways *^12^*. Deubiquitinating enzymes (DUBs) reverse these ubiquitin modifications by removing ubiquitin from target proteins and by disassembling polyubiquitin chains *^13, 14^*. The opposing actions of ubiquitination and deubiquitination are essential for regulating ubiquitin concentrations in eukaryotic cells.

DUBs must be able to navigate the intricacies of the ubiquitin system – differentiating mono- from poly-ubiquitinated species and selectively cleaving one linkage type over another. Humans have ~100 DUBs, which are classified into seven evolutionarily conserved families: the cysteine proteases, ovarian tumor proteases (OTU), ubiquitin C-terminal hydrolases (UCH), ubiquitin specific proteases (USP), Josephins (MJD), MINDY, and ZUP1; and the zinc-dependent Jab/MPN (JAMM) metalloproteases *^13^*. Many DUBs are found in complex with other proteins *^15^*, which in some cases has been shown to modulate DUB activity *^16^*. A mass spectrometry screen of basal isopeptidase activity for 42 DUBs revealed that the majority of these enzymes possess little to no catalytic activity in their apo form *^17^*, raising the possibility that many of these DUBs rely on partner proteins to increase their activity.

OTUB1 is an OTU class DUB that is highly expressed in mammalian cells *^18, 19^* and is highly specific for cleaving K48-linked polyubiquitin chains *^20, 21^*. The deubiquitinating activity of OTUB1 is implicated in removing polyubiquitin chains from, and thereby stabilizing, transcription factors such as FOXM1 *^22, 23^* and ERα *^24^*, as well as E3 ligases such as c-IAP1 *^25^*, TRAF3 and TRAF6 *^26^*, and GRAIL *^27^*. As a result, the DUB activity of OTUB1 regulates transcription, MAPK signaling, virus-triggered type I interferon pathways, and T-cell anergy.

A unique characteristic of OTUB1 is that it binds to a subset of E2 enzymes and inhibits ubiquitin transfer in a manner that is independent of its deubiquitinating activity *^28–31^*. This unusual non-catalytic activity was first reported in studies of the DNA double strand break response, where it was shown that OTUB1 limits accumulation of K63-linked chains at DNA double-strand breaks *^28^*. OTUB1 inhibits synthesis of K63-linked chains by the E2, UBE2N, by binding to the E2~Ub thioester, thus preventing ubiquitin transfer *^28^*. Mass spectrometry-proteomic studies have revealed that OTUB1 interacts with six other E2 enzymes in cells: UBE2D1 (UbcH5a), UBE2D2 (UbcH5b), UBE2D3 (UbcH5c), UBE2E1 (UbcH6), UBE2E2 (UbcH8), and UBE2E3 (UbcH9 or UbcM2) *^15, 28^*. It has been reported that OTUB1 non-catalytically inhibits UBE2D proteins from ubiquitinating p53 *^32, 33^* and SMAD2/3, regulators of TGFβ signaling *^34^*. A recent study showed that OTUB1 also prevents autoubiquitylation of UBE2E1, thereby rescuing the E2 from proteasomal degradation *^35^*. The biological roles of other E2 complexes with OTUB1 remain to be explored.

Structural studies of OTUB1 have provided insights into how this DUB inhibits E2 enzymes. Like other members of the OTU family *^36^*, OTUB1 contains two ubiquitin binding sites, proximal and distal, which bind two ubiquitin monomers, while the K48 isopeptide linkage is positioned near the catalytic cysteine **(Figure 1)***^30, 31, 37^*. A flexible N-terminal ubiquitin-binding helix that forms part of the proximal site is largely disordered in the apo enzyme **(Figure 1a)***^20, 31^* but forms an α-helix when ubiquitin is bound *^31^*. When OTUB1 forms an inhibitory complex with the E2~Ub thioester **(Figure 1d)**, the E2 is oriented such that the conjugated ubiquitin binds to the proximal site in OTUB1*^30, 31^*. Binding of the thioester-linked ubiquitin to the proximal site is stimulated when a second mono-ubiquitin binds in the distal site, thus allosterically increasing the affinity of OTUB1 for the E2-linked ubiquitin in the proximal site *^21, 31^* **(Figure 1b-c)**. The orientation of the proximal E2-linked ubiquitin and distal mono-ubiquitin mimics the expected binding of K48-linked diubiquitin **(Figure 1c)***^30, 31, 37^*.

**Figure 1.**
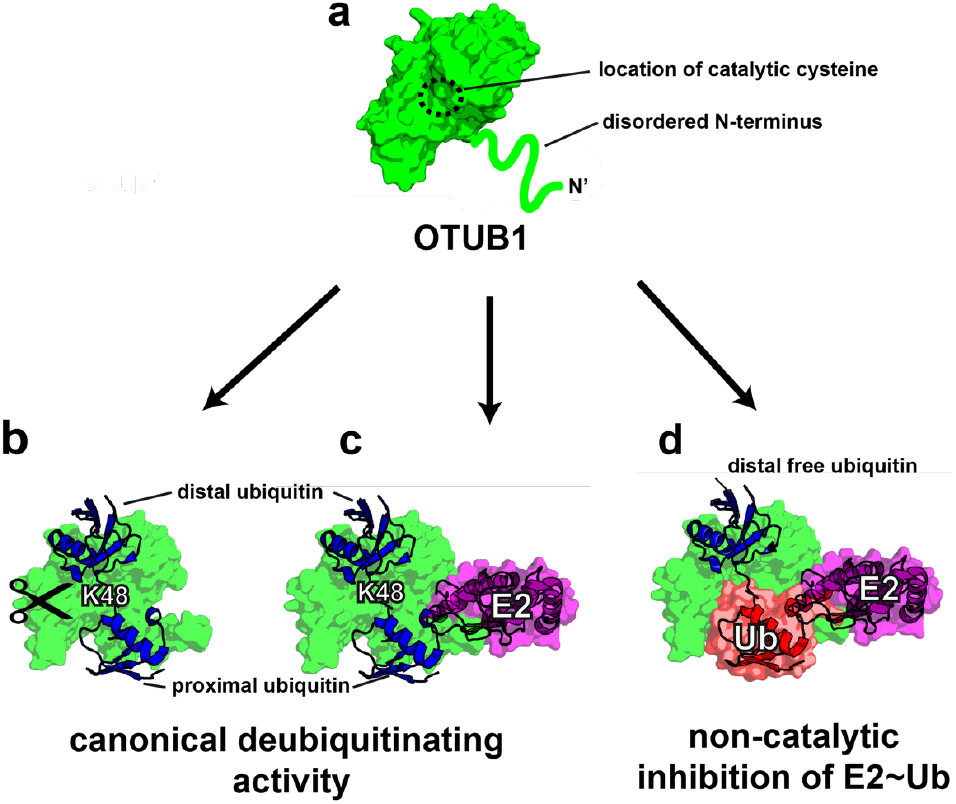
Consequences of OTUB1 binding to its E2 enzyme partners. (a) Schematic of apo-OTUB1 with its disordered N-terminal residues in the absence of substrate (PDB ID 2ZFY). (b) Model of OTUB1 bound to K48 diubiquitin based on PDB: 4LDT structure (OTUB1~Ub aldehyde bound to a charged UBE2D2). The OTUB1 N-terminus is ordered and contacts the proximal ubiquitin. (c) Model of OTUB1 bound to both K48 diubiquitin and an E2 enzyme. (d) Inhibited complex, OTUB1:E2~Ub bound to free Ub on the distal binding site. In red is the conjugated ubiquitin of the charged E2 which binds to the proximal site.

The ability of select E2 enzymes to bind OTUB1 also impacts the isopeptidase activity of the DUB on K48-linked polyubiquitin. An *in vitro* study showed that binding of an uncharged E2 enzyme to OTUB1 can stimulate cleavage of polyubiquitin by stabilizing N-terminal ubiquitin-binding helix to fold which then comprises the proximal ubiquitin-binding site *^37^*. A comparison of the effect of several different E2 enzymes on OTUB1 DUB activity showed that the UBE2D enzymes and UBE2E1 robustly stimulate OTUB1, although UBE2N only weakly enhanced cleavage of K48 diubiquitin under the conditions tested *^37^*. Thus, the same E2 enzymes that are inhibited by OTUB1 when the E2 is charged with ubiquitin can stimulate OTUB1 when uncharged. Indeed, the charged E2~Ub can also be said to regulate OTUB1 activity, since OTUB1 is unable to cleave polyubiquitin when it forms a repressive complex with E2~Ub and Ub. These observations suggested a potential network of cross-regulation in cells in which OTUB1:E2 complexes have different enzymatic activity depending on whether or not the E2 enzyme is charged with ubiquitin *^37^*.

The contribution of E2 binding to OTUB1 DUB activity *in vivo*, as well as the relative balance between active and repressed OTUB1 E2 complexes, is unknown. The K_M_ of OTUB1 for K48 diubiquitin has been reported to be in the range of 84 - 120 μM *^20, 21, 37^*. These K_M_ values are far above the physiological concentration of polyubiquitin in cells, which varies depending on the cell type, but is estimated to be at a concentration below 10 μM for all types of chains *^38, 47^*. This therefore suggests that OTUB1 alone is not likely to cleave polyubiquitin *in vivo*. Binding of UBE2D2 to OTUB1, however, lowers the K_M_ for of OTUB1 K48 chains by more than an order of magnitude to within range of physiological substrate concentrations *^37^*. However, it is not known whether the remaining OTUB1-interacting E2s similarly regulate DUB activity over a plausible biological concentration. The relative balance of activities *in vivo* depends on the cellular concentrations of the proteins and the relative kinetics and thermodynamics of these interactions, which have remained understudied.

We report here a kinetic and thermodynamic characterization of OTUB1 binding to its E2 partners and the effect of complex formation on OTUB1 affinity for, and cleavage of, K48 polyubiquitin. To fully examine the cross-regulatory network, we also quantified the inhibition of OTUB1 by its E2 partners under similar conditions. Together, these measurements allowed us to elucidate the relative thermodynamic balance between E2 inhibition and DUB stimulation, and to show that DUB stimulation occurs over a range of enzyme concentrations that correspond to those measured in cells *^18, 19^*. These parameters provide a framework for investigating the diverse *in vivo* roles of OTUB1-E2 complexes.

## Materials and Methods

### Cloning and mutagenesis

Plasmids used for *E. coli* expression of human Uba1 (E1) *^39^*, RNF4 *^40^*, all E2 enzymes (UBE2D1, UBE2D2, UBE2D3, UBE2N, UBE2E1, and Cdc34 (UBE2R1q) *^37^*, and *Saccharomyces cerevisia*e Ubiquitin C-terminal hydrolase-1 (yUH1) *^41^* were previously described. UBE2E2 and UBE2E3 were cloned from a human cDNA library (Clontech) into the pET-SUMO2 vector (Addgene) that contains an N-terminal His_6_-tag followed by a SENP2 protease site. OTUB1 wild-type was cloned into pProEx-c vector (Life Technologies) containing a N-terminal His_6_-TEV protease site as previously described *^21^*. The catalytic mutant, OTUB1 (C91S), was made by site-directed mutagenesis. Ubiquitin (Ub) wild-type and Ub G77D were cloned into pET3a plasmids as previously described *^21^*. For Ub K(48/63)R mutants, Quikchange cloning (Clontech) was used to introduce both mutations into the pET3a-Ub wild-type plasmid. All clones were transformed into XL1-Blue cells (Stratagene).

### Protein expression and purification

All proteins were expressed in *E.coli* Rosetta-2 DE3 cells (EMD Millipore, Merck KGaA, Darmstadt, Germany). Cells were transformed via heat shock (42°C for 35 secs) and plated on Luria-Bertani broth (LB) agar plates with 34 mg/mL chloramphenicol and 100 mg/mL carbenicillin. A small starter growth of 50 mL was initiated by picking 2-5 Rosetta-2 DE3 colonies and placing into LB media with 34 mg/mL chloramphenicol and 100 mg/mL carbenicillin antibiotics. Starter cultures were grown at 37°C shaking at 250 rpm overnight. For yUH1, E1, all E2s (UBE2D1, UBE2D2, UBE2D3, UBE2N, UBE2E1, and Cdc34A), and all OTUB1 variants (wild-type and C91S) large-scale cultures (1L) were composed of M9ZB media (1x M9 salt mix, 77 mM NaCl, 10 g/L casamino acids, 2 mM MgSO_4_, 0.5% glycerol, and 0.5x Metal mix) that were inoculated with 1% (v:v) overnight saturated starter cultures containing 34 mg/mL chloramphenicol and 100 g/mL carbenicillin. Cultures were grown at 37°C to an of O.D_600_ ~1.5, induced by the addition of 1 mM isopropyl βD-1-thiogalactopyranoside (IPTG) and incubated overnight (~16 hrs) at 16°C. For all monoubiquitin variants (wild-type, Ub-G77D, and Ub-K48R/63R) a 50 mL starter growth of LB containing: 2-5 Rosetta-2 DE3 colonies, 34 mg/mL chloramphenicol and 100 mg/mL carbenicillin antibiotics, were grown at 37°C shaking at 250 rpm until the O.D_600_ reached 0.4-0.6 and was placed in 4°C overnight to halt growth. Large scale cultures (1L) of 2xYT media (Millipore-Sigma) were made by adding 1% (v:v) O.D_600_ 0.4-0.6 starter culture, 34 mg/mL chloramphenicol, and 100 g/mL carbenicillin. These cultures were grown at 37°C to an of O.D_600_ 0.8-1.0, induced by the addition of 1 mM isopropyl βD-1-thiogalactopyranoside (IPTG) and incubated overnight (~16 hrs) at 16°C. Cells were harvested by centrifuging at 4000 rpm at 4°C then immediately lysed with a Microfluidizer (Microfluidics).

yUH1 *^41^* and E1 *^39^* were purified as previously described. Wild type ubiquitin, Ub-G77D, and Ub-K48R/63R proteins were purified as previously described *^21^*. All E2 enzymes (UBE2D1-3, UBE2N, UBE2E1-3 and Cdc34A) and OTUB1 proteins (wild-type and C91S) were purified by resuspending pelleted cells in lysis buffer (20 mM HEPES pH 7.3, 300 mM NaCl, 25 mM imidazole, 2 mM β-mercaptoethanol (BME)). Phenyl-methyl sulphonyl fluoride (PMSF), 1mM, was added to cells before lysis with a microfluidizer (Microfluidics). The lysate was centrifuged (12,500 rpm at 4°C for 25 mins) and purified by affinity chromatography on a 5mL HisTrap HP column (GE Healthcare Life Sciences). Proteins of interest were eluted using a linear gradient of 0 - 250 mM imidazole over 10 column volumes. To cleave the His-tag, 10 mM His_6x_-SENP2 was added to the eluted fraction at a v:v ratio of 1:100, and dialyzed overnight at 4°C in lysis buffer. A second round of HisTrap purification was used to separate the cleaved protein from the tag and from His_6x_-SENP2. The flow-through was collected and further purified as described below.

After removing the His_6x_-tag, all E2 enzymes (UBE2D1, UBE2D2, UBE2D3, UBE2N, and Cdc34A) except UBE2E1-3 were purified further by size exclusion chromatography. Fractions were concentrated to 2 mL and run on a HiLoad 16/60 Superdex 75 column with buffer: 50 mM HEPES pH 7.5, 150 mM NaCl, and 0.5 mM Tris(2-carboxyethyl)phosphene (TCEP)-HCl. Eluted fractions were then concentrated down to 4-10 mg/mL, flash-frozen in liquid nitrogen, and stored at −80°C.

After cleaving the His_6x_-tag overnight, the UBE2E proteins were passed through a HisTrap HP column and flow-through fractions were collected and dialyzed overnight into 25 mM sodium phosphate buffer pH 7.4, 25 mM NaCl, and 7.5 mM BME at 4°C. UbE2E1-3 were further purified using cation exchange chromatography (SP HP 5 mL GE Lifesciences) and eluted at 100 mM NaCl. Pure fractions were dialyzed overnight at 4°C in 25 mM Tris buffer pH 8, 150 mM NaCl and 7.5 mM BME, concentrated to 4-10mg/mL, and stored at −80°C.

After the His_6x_-tag was removed, OTUB1 and its catalytic mutant (C91S) were further purified by gel filtration chromatography on a HiLoad 16/60 Superdex 75 (GE Healthcare Life Sciences) column equilibrated in 25 mM HEPES pH 7.6, 150 mM NaCl, and 0.5 mM tris(2-carboxyethyl) phosphine (TCEP). Purified OTUB1 was concentrated to 4-10 mg/mL, frozen and stored at −80°C until use.

### Synthesis of K48-linked diubiquitin

Reaction mixes containing 7 μM Cdc34A, 700 μM Ub K(48/63)R and 500 μM Ub G77D were prepared in a buffer of 50 mM Tris pH 7.6, 10 mM MgCl_2_, and 1 mM Dithiothreitol (DTT). Reactions were initiated with the addition of 0.1 μM E1 and 10 mM ATP pH 7.5, then incubated for 18 hrs at 37°C. In order to remove the C-terminal D77 residue from the C-terminus of the proximal ubiquitin, 10 μM of the DUB, yUH1, was added to the reaction mix, along with 1 mM DTT and 10 mM EDTA, and incubated at 37°C for an additional 3 hrs *^42^*. The mixture was then diluted 10-fold with 50 mM ammonium acetate pH 4.5, 10 mM NaCl, and 1 mM DTT, and filtered to remove any aggregated proteins. Reactions were purified using and fractions containing K48 diubiquitin were separated from unreacted monoubiquitin using cation exchange chromatography with a linear gradient with 50 mM ammonium acetate pH 4.5, 10-600 mM NaCl, and 1 mM DTT. Fractions containing pure K48 diubiquitin as judged by SDS-PAGE were pooled, concentrated to 10 mg/mL and stored at −80°C.

### Isothermal Titration Calorimetry (ITC)

All ITC experiments were performed on a MicroCal ITC_200_ system (Malvern) at 25°C. The syringe (70 μL) titrated protein into a 300 μL sample cell. Each ITC experiment contained 19 injections of 2 μL each for a duration of 0.8 sec, with 150 sec between injections. All proteins were first dialyzed in a buffer of 25 mM HEPES, pH 7.5, 150 mM NaCl, and 0.5 mM TCEP-HCl. For measurements of interactions between E2 and OTUB1 (C91S), the sample cell contained 150 μM of E2 and the syringe 1.5 mM OTUB1. For OTUB1 (C91S) interactions with K48 diUb, 1.5 mM K48 diubiquitin was titrated into 150 μM of OTUB1. For experiments testing the binding of K48 diubiquitin to OTUB1 (C91S) at saturating concentrations of E2, 3 mM K48 diubiquitin was titrated into 150 μM OTUB1 (C91S) with a constant 150 μM of E2 present in both the syringe and the cell. Heat generated due to dilution of the titrants was subtracted for baseline correction. The baseline-corrected data were analyzed with MicroCal Origin Ver. 7.0 software.

### Kinetic assays of OTUB1 deubiquitinating activity

The K48 diubiquitin substrate used for all assays was internally quenched fluorescent (IQF) substrate no. 5 (LifeSensors) FRET-K48 diubiquitin, which contains TAMRA conjugated to one ubiquitin and a quencher to the other. Reaction volumes of 30 μL were maintained at 30°C on a POLARStar Omega plate reader (BMG LABTECH) that measured TAMRA fluorescence (ex. 544 nm; em. 590 nm) every 5 secs over 30 mins. To generate a standard curve, FRET-K48 diubiquitin at concentrations ranging from 25 - 500 nM was fully digested by incubation with 50 nM OTUB1 for 1 hr at 30°C. Gain was adjusted to 1900 based on the fluorescence from the 500 nM FRET-K48 diubiquitin reaction. The standard curve was obtained by plotting fluorescence (AU) as a function of concentration of the cleaved FRET-K48 diubiquitin. The data were then fitted to a linear equation to obtain the slope in units of AU •μM^−1^. This slope was used to convert AU into a measurement of concentration of cleaved diubiquitin.

For the fold stimulation assay, reactions (30 μL) containing 400 nM of FRET-K48 diubiquitin in a buffer of 20 mM HEPES, pH 7.5, 150 mM NaCl, 1 mM DTT, and 0.01% BSA were assayed at 30°C. Reactions performed in the absence and the presence of 10 μM E2 enzyme and were initiated by the addition of 50 nM OTUB1. Data were analyzed in GraphPad Prism. Experiments were done in triplicate.

To determine EC_50_ values for OTUB1 cleavage of K48 diubiquitin, reactions were measured at 30°C in buffer containing 20 mM HEPES, pH 7.5, 150 mM NaCl, 1 mM DTT, 0.01% BSA, and 400 nM of FRET-K48 diubiquitin. Reactions (30 μL) contained specified amounts of E2 enzyme and were initiated by addition of 50 nM OTUB1. The initial rate of Lys48 diubiquitin cleavage was determined from the slope of the linear region of the fluorescent curves. Data were analyzed in GraphPad Prism. Experiments were done in triplicate.

Steady-state enzyme kinetic assays were performed at 30°C in buffer containing 20 mM HEPES, pH 7.5, 150 mM NaCl, 1 mM DTT, 0.01% BSA, in a reaction volume of 50 μL. Reactions in the presence and absence of 10 μM E2 were initiated by addition of 50 nM OTUB1. For all experiments, the concentration of unlabeled K48-linked diubiquitin was increased in the presence of a constant 400 nM FRET-K48 diubiquitin. After each incubation, the actual concentration of ubiquitin cleaved was obtained by correcting for the proportion of FRET-labeled versus unlabeled K48 diubiquitin. The transformed data were plotted as a function of time and fit to a line where initial velocity conditions were satisfied, typically within the first three minutes. Initial rates were measured in triplicate, normalized to the enzyme concentration, plotted as a function of substrate concentration, and the resulting curve fit to the Michaelis-Menten equation using non-linear least squares regression implemented in GraphPad Prism 5 (GraphPad Software).

### Inhibition assays

Inhibition of E2 ubiquitin-conjugating activity was assayed with an SDS-PAGE gel-based assay. Each initial reaction mix contained 2 μM E2, 2 μM RNF4 (the E3 ligase), and 50 μM ubiquitin in the presence and absence of specified concentrations of OTUB1 (C91S) in reaction buffer containing 50 mM HEPES pH 7.5, 300 mM NaCl, 10mM MgCl_2_, 0.5 mM DTT, and 0.005 % Tween20 at 37°C. Reactions (10 μL) were initiated by addition of 5 mM ATP, pH 7.5 and 0.1 μM E1 and quenched at the specified time points by addition of SDS-PAGE sample buffer containing BME. Samples were electrophoresed on 4–12% gradient polyacrylamide Bis-Tris Criterion XT gels (Bio-Rad) and were either stained with Coomassie Brilliant Blue (Bio-Rad) or transferred to a PVDF membrane (Bio-Rad) for Western blotting using a Bio-Rad Turbo Transfer system. All blots were blocked with 5% BioRad blotting grade blocker in Tris-buffered saline and Tween20 solution (TBST) for 1 hr at room temperature. Membranes were then washed with TBST, rocking at room temperature for 10 mins. Blots were then incubated overnight at 4°C with primary antibody **(Table S1)** diluted in 2% BSA, 0.02% sodium azide in phosphate buffered saline (PBS). Blots were washed again three times with TBST for 10 min intervals before adding secondary antibody **(Table S2)**, diluted in 5% BioRad blotting grade blocker, and shaken at room temperature for 1 hr. After another round of washes with TBST, 1:1 BioRad Clarity ECL reagent was used to visualize proteins of interest.

## Results

### Differences in stimulation of OTUB1 are not correlated with E2 binding affinity

The ability of a subset of E2s to stimulate OTUB1 has previously been reported *^37^* but the consequences of UBE2E2 and UBE2E3 binding on OTUB1 DUB activity had not been examined. We therefore compared the ability of UBE2E2 and UBE2E3 to stimulate the DUB activity of OTUB1 to that of the other OTUB1 interacting E2 partners: UBE2E1, UBE2N, and the three UBE2D enzymes. Using a Förster resonance energy transfer (FRET) deubiquitination assay (see Methods), we found that UBE2E2 and UBE2E3 only modestly enhanced the DUB activity of OTUB1 **(Figure 2a)**. As compared to UBE2D2, a robust stimulator that enhanced DUB activity by 25-fold, UBE2E2 and UBE2E3 only stimulated OTUB1 by 5-fold, which was the weakest for all E2s studied.

**Figure 2.**
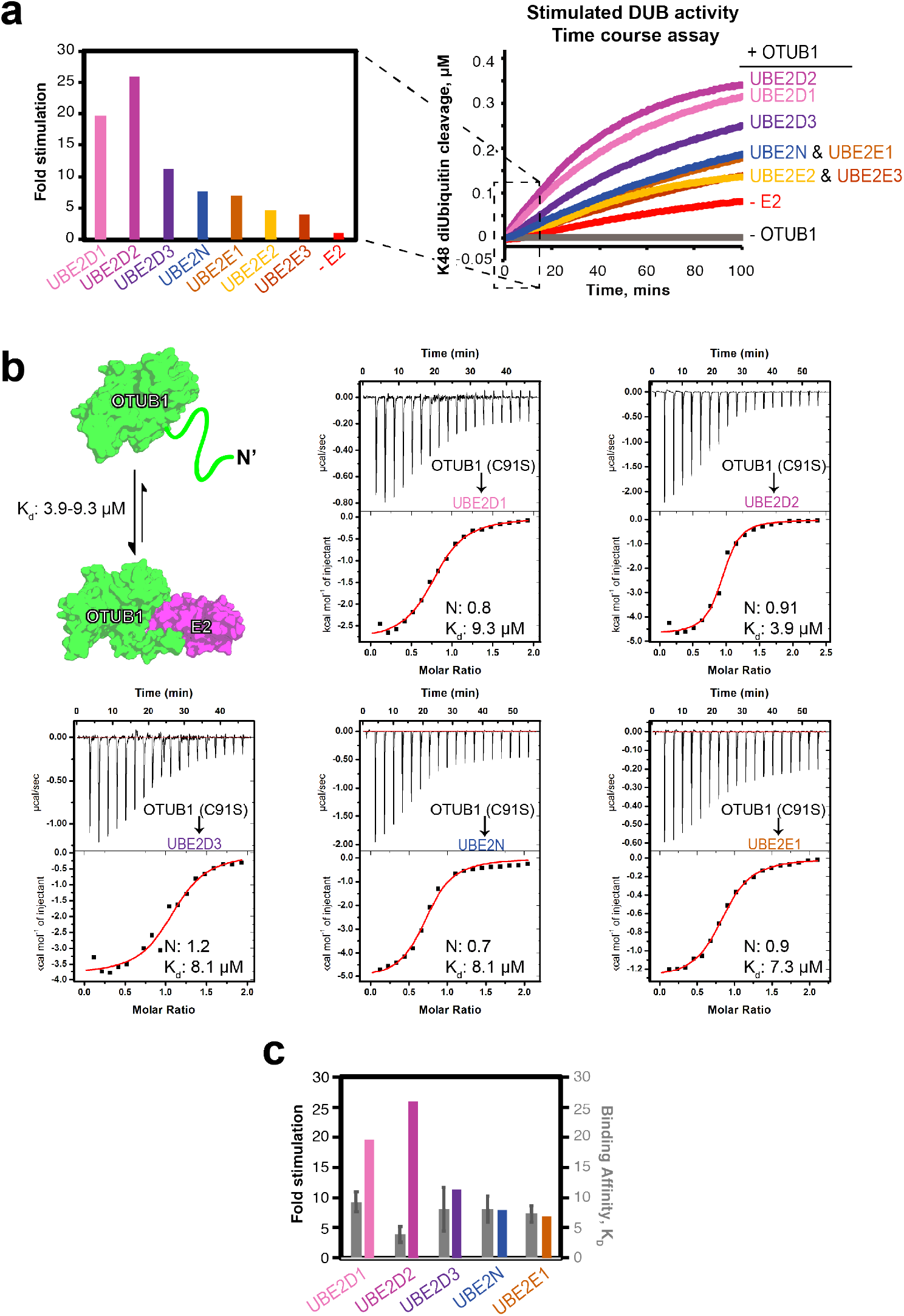
E2 stimulation of OTUB1 and E2-OTUB1 affinity. (a) Fold stimulation of OTUB1 (50 nM) activity by the indicated E2 enzymes (10 μM) (left) determined from initial reaction rates in the FRET-based time course assay depicting cleavage of internally quenched fluorescent K48 diUbiquitin (400 nM) (right). (b) Binding schematic and ITC experiments for OTUB1 binding to the indicated E2 partner. OTUB1 (1.5 mM) was titrated into the corresponding E2 (150 μM), contained within the cell. (c) Side-by-side comparison of *K*_*d*_ values measured in (b, colored grey and right axis) with the fold stimulation determined in (a, multicolored and left axis) for the indicated E2s.

Formation of both active and repressive OTUB1-E2 complexes are governed, in part, by the intrinsic affinity of OTUB1 for each of its E2 partners. To see if differences in relative stimulation of DUB activity can be attributed to variations in binding affinities, we determined equilibrium dissociation constants (K_d_) for OTUB1 binding to UBE2D1, UBE2D2, UBE2D3, UBE2N, and UBE2E1 using isothermal titration calorimetry (ITC). For these experiments, we used the catalytically inactive mutant, OTUB1 (C91S). The ITC experiments showed that the affinities between each E2 and OTUB1 were similar, with K_d_ values ranging from 3.9 – 9.3 μM **(Figure 2b; thermodynamic parameters listed in Table S3)**. The K_d_ for binding of UBE2N to

OTUB1 was 8.1 μM, which is consistent with previous studies that measured a K_d_ of 7.04 μM using fluorescence polarization *^31^* and K_d_ of 8.9 μM measured by surface plasmon resonance using a GST-OTUB1 fusion *^29^*. In order to verify that the OTUB1 C91S mutation did not affect binding to E2 enzymes, we measured binding of a subset of E2s (UBE2D1 and UBE2D3) to wild-type OTUB1 found no significant difference in K_d_ values **(Figure S1)**. We were unable to measure the K_d_ for the interactions between OTUB1 and UBE2E2 or UBE2E3, as the two proteins precipitated in the ITC cell. Taken together, the small differences in K_d_ observed between OTUB1 and E2 partners do not explain the large differences in the ability of different E2 enzymes to stimulate OTUB1 DUB activity **(Figure 2c)**.

UBE2G2 was not originally reported as an OTUB1 binding partner *^28^* but was subsequently identified as a partner by affinity capture mass spectrometry *^43^*. However, we saw no interaction between OTUB1 and UBE2G2 by ITC **(Figure S2a)**. The thermogram is similar to that of CDC34 **(Figure S2b)**, which is not thought to bind OTUB1 and does not stimulate its activity *^37^*.

### EC_50_s for E2 inhibition of OTUB1 match intracellular concentrations

To further characterize how E2 binding to OTUB1 effects DUB activity, we assayed OTUB1 activity as a function of E2 concentration to determine the half-maximal effective E2 concentration (EC_50_) for stimulation of OTUB1. Using a FRET-based assay, we measured rates of OTUB1 isopeptidase activity over increasing log_10_ concentrations of E2 (0.001 nM – 100 μM). We found that the E2 concentration needed to elicit a stimulatory response ranged from 0.9 to 16.1 μM **(Figure 3)**. Of the seven E2s studied, the UBE2N EC_50_ of 0.8 μM was the lowest measured. UBE2E2 and UBE2E3 had the highest EC_50_ values of at least 12.1 and 16.1 μM, respectively, although these values are likely an underestimate as it was not possible to reach a saturating concentration for stimulation **(Figure 3)**. The effective concentrations for the UBE2D family of enzymes were in the range of 1.6 – 4.3 μM. The EC_50_ values determined for UBE2N and UBE2D3 match their intracellular concentrations in mouse embryonic fibroblasts *^19^*, which are 1.3 μM for UBE2N and 1.7 μM for UBE2D3. Taken together, our calculated EC_50_ values suggest that these E2 partners can regulate OTUB1’s activity over a physiological range of concentrations.

**Figure 3.**
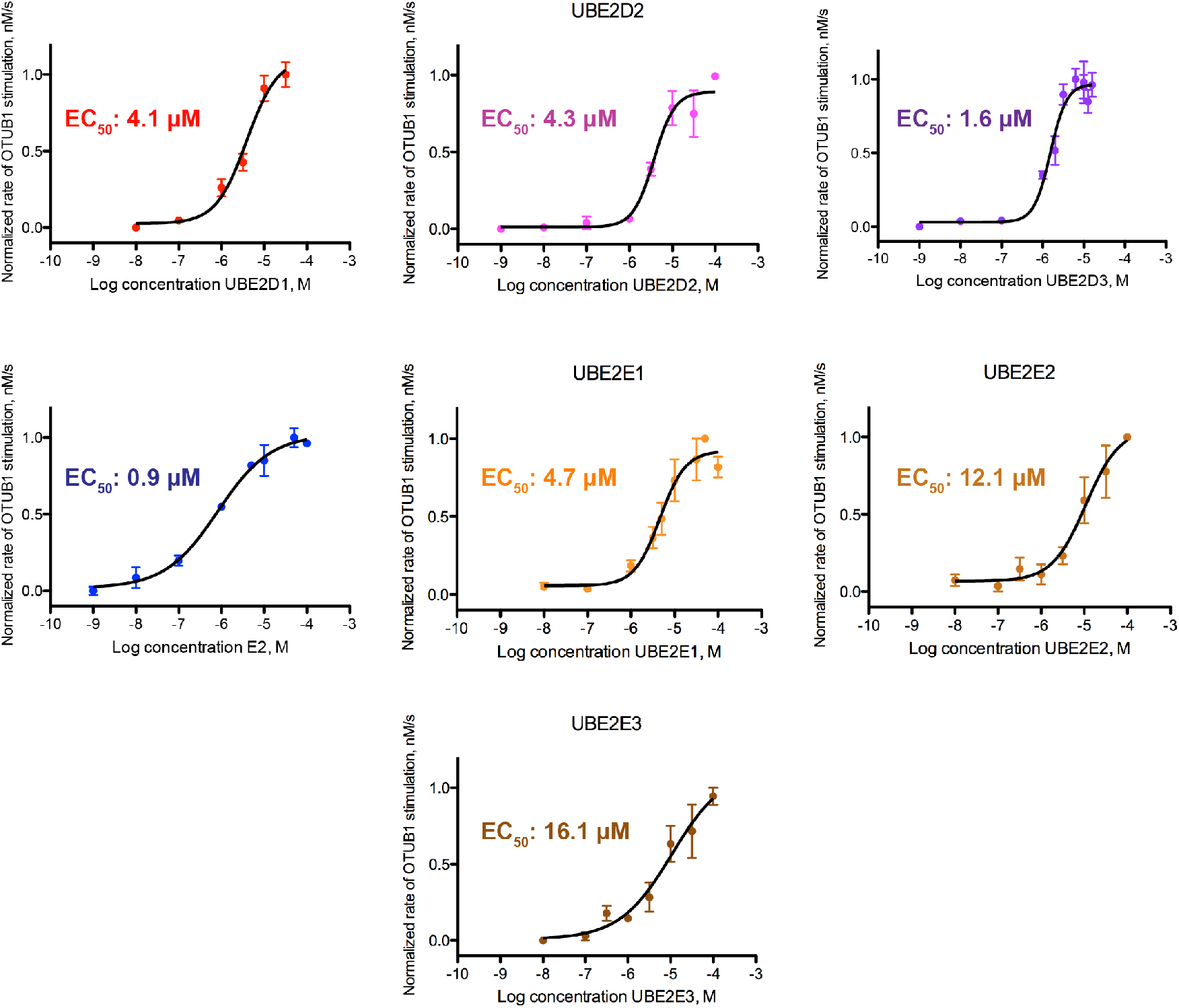
Effective concentration (EC_50_) of E2 stimulation of OTUB1 activity. Cleavage of FRET-K48 diUbiquitin (400 nM) by OTUB1 (50 nM) was assayed with increasing concentrations of E2 enzymes (1 pM to ~50 μM). Reactions were assayed in triplicate.

### Binding an E2 partner raises affinity of OTUB1 for K48 diubiquitin

A previous study *^37^* reported that the binding of UBE2D2 lowered the K_M_ of OTUB1 for K48 diubiquitin by 34-fold, from 120 μM to 3.4 μM, with no significant change in k_cat_. The effect of other E2 partners on OTUB1 kinetics, however, had not been studied. We therefore determined the effects of saturating concentrations of UBE2D1, UBE2D3, UBE2N, and UBE2E1 **(Figures 4 and S3**) on the kinetics of OTUB1 DUB activity. UBE2D2 was included to provide an internal comparison, since previous kinetic parameters were determined with different methods *^37^*. In the absence of an E2, the K_M_ of OTUB1 for K48 diubiquitin was 102 μM and the *k*_*cat*_ was 0.03 s^−1^ **(Figure 4)**. These values are, within error, comparable to previously reported *K*_*M*_ values of 78 μM *^21^* and 120 μM *^37^*, and a k_cat_ of 0.034 s^−1^ *^37^*. We found that the UBE2D isoforms lowered the *K*_*M*_ of OTUB1 for K48 diubiquitin to the greatest extent, with K_M_ values of 11 μM in the presence of UBE2D1, 6.6 μM in the presence of UBE2D2, and 13 μM in the presence of UBE2D3. The *K*_*M*_ of OTUB1 for substrate in the presence of UBE2N was 24.1 μM, consistent with the lower levels of stimulation observed in our time course assays **(Figure 2a)**. Stimulation of DUB activity was weakest in the presence of UBE2E1, which lowered the K_M_ less than 2-fold to 64.3 μM. We did not quantify changes in DUB activity in the presence of either UBE2E2 or UBE2E3; however, given their even lower ability to stimulate OTUB1 **(Figure 2a)**, we speculate that they have an even smaller effect on *K*_*M*_ than UBE2E1. Importantly, the presence of an E2 in all cases affected only *K*_*M*_, with little to no change in *k_cat_*. A plot of *k*_*cat*_/*K*_*M*_ versus the fold stimulation observed in **Figure 2a** reveals a linear correlation (Figure 4b), indicating that the increase in isopeptidase activity is entirely the result of changes in *K*_*M*_ and not *k*_*cat*_.

**Figure 4.**
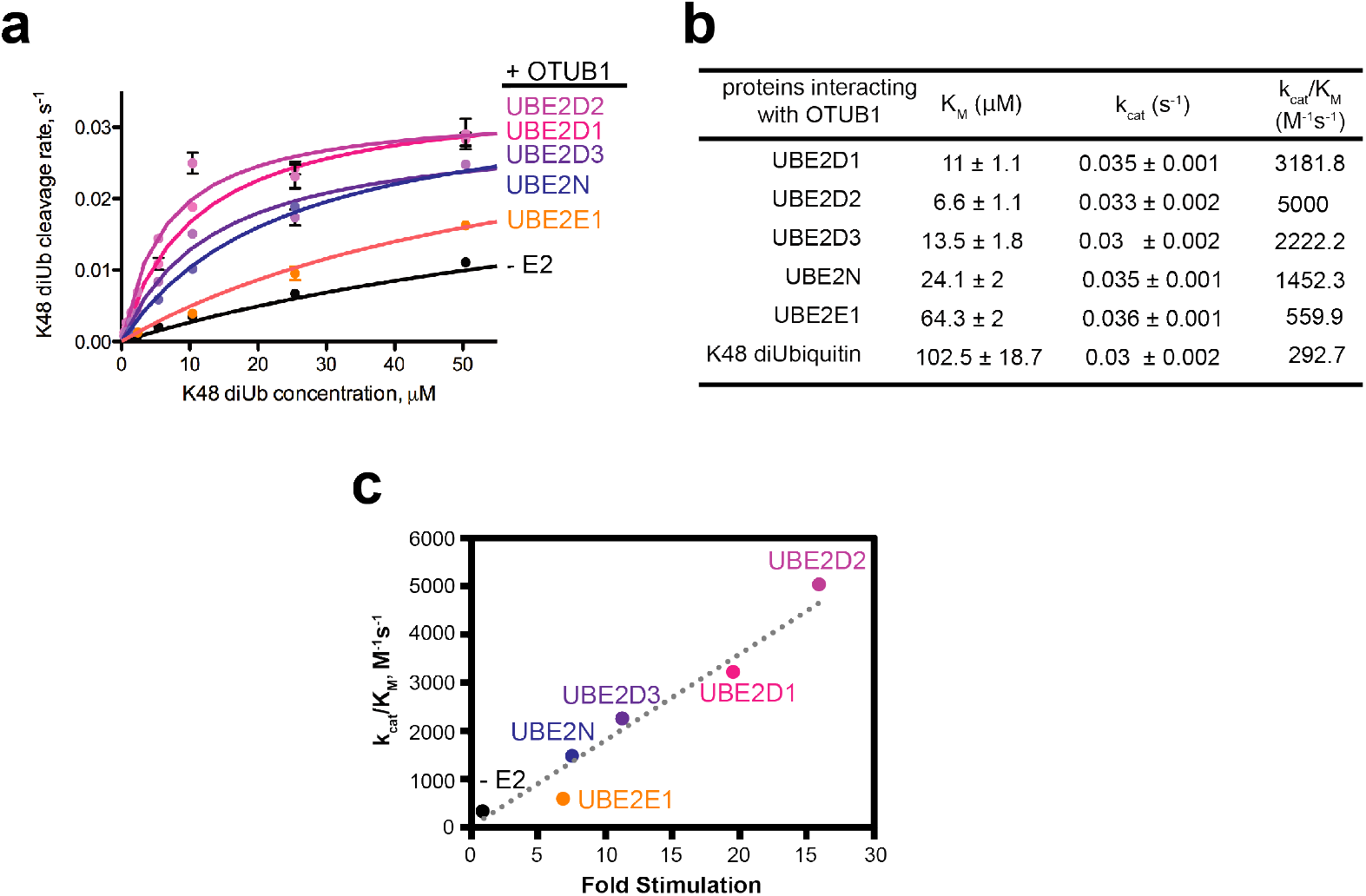
All E2 partners lower the *K_M_* and *K_d_* of OTUB1 for K48 diUbiquitin. (a) Kinetic assay of OTUB1 (50 nM) DUB activity in the presence and absence of 10 μM E2, which is above the K_d_ of all the E2 partners. Reactions were performed in triplicate. The full substrate titration up to 500 μM K48 diUbiquitin can be found in Supplemental Figure S3. (b) Summary table of all determined kinetic parameters: *K*_*M*_, *k*_*cat*_, and the calculated *k*_*cat*_/*K*_*M*_. (c) Correlation between changes in *k_cat_*/*K_M_* measurements to the increase in fold stimulation for isopeptidase activity (from Figure 2a) in the absence and presence of certain E2 enzymes. Dotted line denotes the linear regression with y = 179.3 M^−1^ s^−1^ and R^2^ = 0.95

The effect of E2 enzymes in lowering the *K*_*M*_ of OTUB1 for substrate suggests that the binding of an E2 partner increases the affinity of OTUB1 for K48 diubiquitin. We tested this hypothesis by using ITC to measure the affinity of OTUB1 for K48 diubiquitin in the presence and absence of E2. For these experiments, we used the catalytic mutant of OTUB1, C91S, to eliminate substrate cleavage. In the absence of E2, we found that OTUB1 binds K48 diubiquitin with a K_d_ of 80 μM (**Figure 5a)**, which is within the range of *K*_*M*_ values reported here **(Figure 4b)** and previously *^21, 37^*. To measure the effect of E2 binding on the affinity of OTUB1 for K48 diubiquitin, ITC measurements were performed in the presence of saturating concentrations of E2 (150 μM) in both the cell and the syringe. This approach allowed us to directly measure K48 diubiquitin binding to the OTUB1:E2 complex without dilution of the E2. We first verified that there was no observable binding between E2 enzymes and K48 diubiquitin at the concentrations used **(Figure S4)**. We determined equilibrium dissociation constants for OTUB1 (C91S) binding to K48 diubiquitin in the presence of UBE2D1, UBE2D3, and UBE2N **(Figure 5b)**. The E2 increased the affinity of OTUB1 for K48 diubiquitin, as reflected in a decrease in K_d_ from 84 μM with no E2 to 12 μM in the presence of UBE2D1, 13.2 μM in the presence of UBE2D3, and 22.3 μM in the presence of UBE2N. These K_d_ values are comparable to the *K*_*M*_ values of OTUB1 in the presence of these E2 partners **(Figure 5e)**, indicating that all E2 enzymes stimulate OTUB1 by increasing the affinity of the DUB for its K48 diubiquitin substrate. This finding directly supports the model that E2 enzymes increase the affinity of ubiquitin for the proximal site by promoting folding of the ubiquitin-binding helix, thereby pre-paying the energetic cost of helix formation *^37^*.

**Figure 5.**
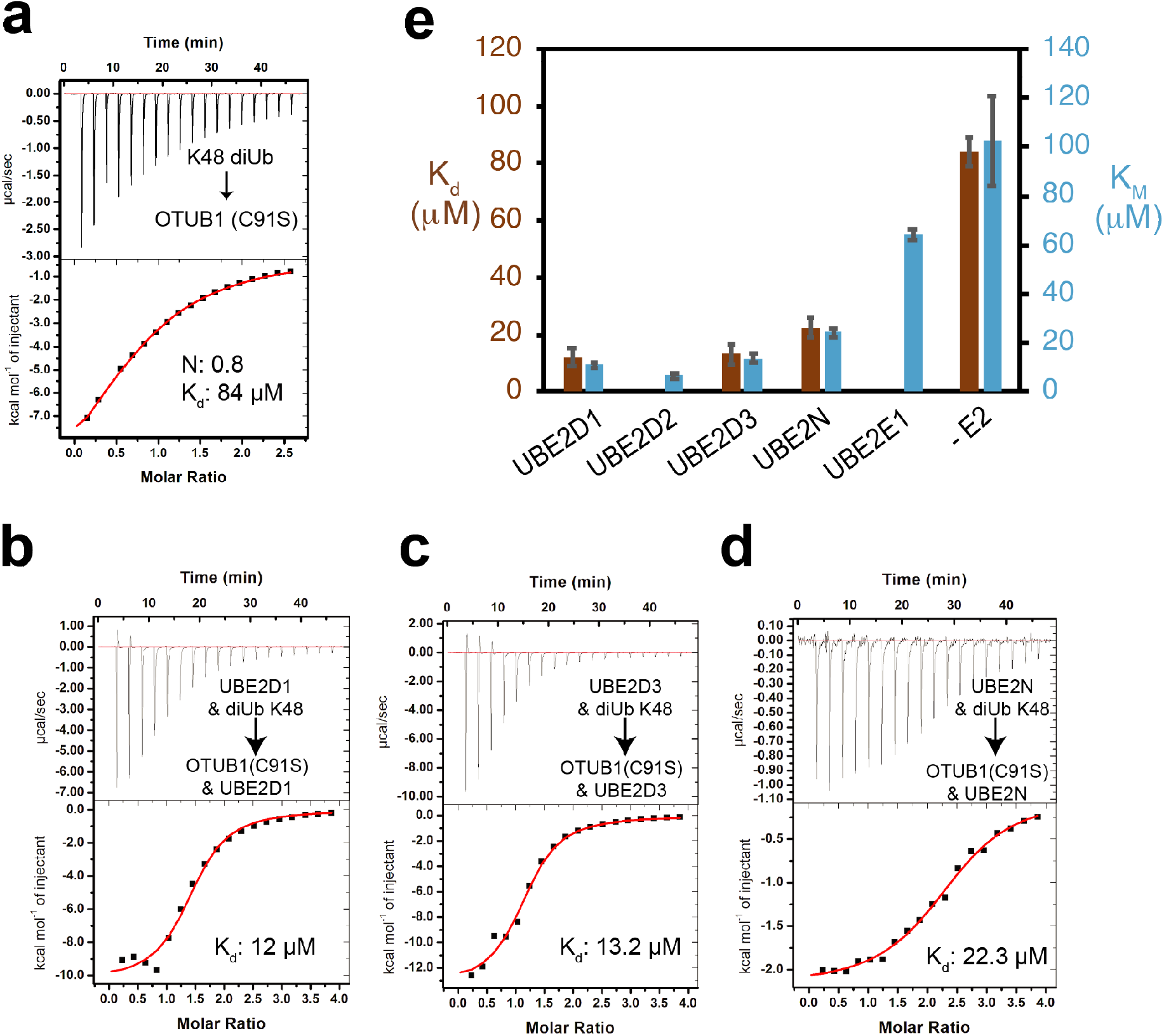
Binding of E2 to OTUB1 raises the affinity of OTUB1 for K48 diUbiquitin. (a) Binding of K48 diUbiquitin, 3 mM, to OTUB1(C91S), 150 μM. (b) Similar experiment as in (a) but in the presence of 150 μM UBE2D1, in both cell and syringe. (c) As in (b), with the E2, UBE2E3. (d) As in (b) with the E2, UBE2N. (e) Bar graph comparing K_d_ values in (a) – (d) with K_M_ values from **Figure 4**.

### OTUB1 inhibits auto-monoubiquitylation of UBE2E1 and UBE2E2 but not UBE2E3

UBE2E1, UBE2E2, and UBE2E3 differ from the other OTUB1-interacting E2s in that they are Class III E2s, whose conserved UBC domains are preceded by an N-terminus that is predicted to be disordered *^44^*. UBE2E enzymes have been shown to primarily mono-ubiquitinate E3 ligases *^45^* and autoubiquitinate lysines in their own N-terminal extension *^46^*. In the case of UBE2E1, autoubiquitylation of a lysine in the N-terminus restricts E2 polyubiquitinating activity *^46^*. It was recently shown that knockdown or knockout of OTUB1 in cells destabilizes UBE2E1 by relieving non-catalytic inhibition of UBE2E1 autoubiquitination, thus causing the E2 to accumulate K48-linked polyubiquitin chains and leading to proteasomal degradation *^35^*. The effect of OTUB1 on the activity of UBE2E2 and UBE2E3 has not been explored.

We assayed the ability of OTUB1 to inhibit autoubiquitylation by UBE2E2 and UBE2E3 in the presence of the E3 ligase, RNF4, which stimulates the activity of all three enzymes. Since the previous study of UBE2E1 *^35^* only assayed inhibition at a single concentration of OTUB1, this E2 was also included in the assays for comparison. We used a gel-based assay and immunoblotting to monitor the ubiquitylating activity of each E2 over time in the presence and absence of an E3 ligase that stimulates UBE2E enzymes, RNF4 *^35, 46^*, as well as with increasing concentrations of OTUB1 **(Figure 6a)**. In the absence of RNF4, all three UBE2E proteins are autoubiquitinated, primarily by attachment of a single ubiquitin and for some, a small amount of higher molecular weight ubiquitin chains **(Figure 6b)**. A small amount of K48-linked diubiquitin is also generated **(Figure 6b)**. As previously reported *^35, 46^*, RNF4 stimulates the activity of all UBE2E enzymes, resulting in polyubiquitinated E2 and a concomitant increase in K48-linked polyubiquitin **(Figure 6b)**, which is likely anchored to either the E2 or to RNF4 *^46^*. As the OTUB1 concentration increased from 0.1 - 50 μM, there was a corresponding decrease in autoubiquitylation of UBE2E enzymes **(Figure 6a)** and K48-linked polyubiquitin synthesis **(Figure 6b)**. The most dramatic reduction in UBE2E activity occurred in the presence of 1-10 μM OTUB1, a similar range over which OTUB1 inhibits the UBE2D isoforms **(Figure S6)**. Unexpectedly, it was not possible to fully inhibit autoubiquitination of UBE2E3 even at concentrations of OTUB1 as high as 50 μM, which is far above physiological levels. Whereas OTUB1 inhibited formation of higher molecular weight autoubiquitinated UBE2E3, monoubiquitinated UBE2E3 persisted and showed little change across all OTUB1 concentrations tested **(Figure 6a**, **right-most panel)**. These results suggest that OTUB1 may serve to primarily inhibit polyubiquitination of UBE2E3, but not autoubiquitylation.

**Figure 6.**
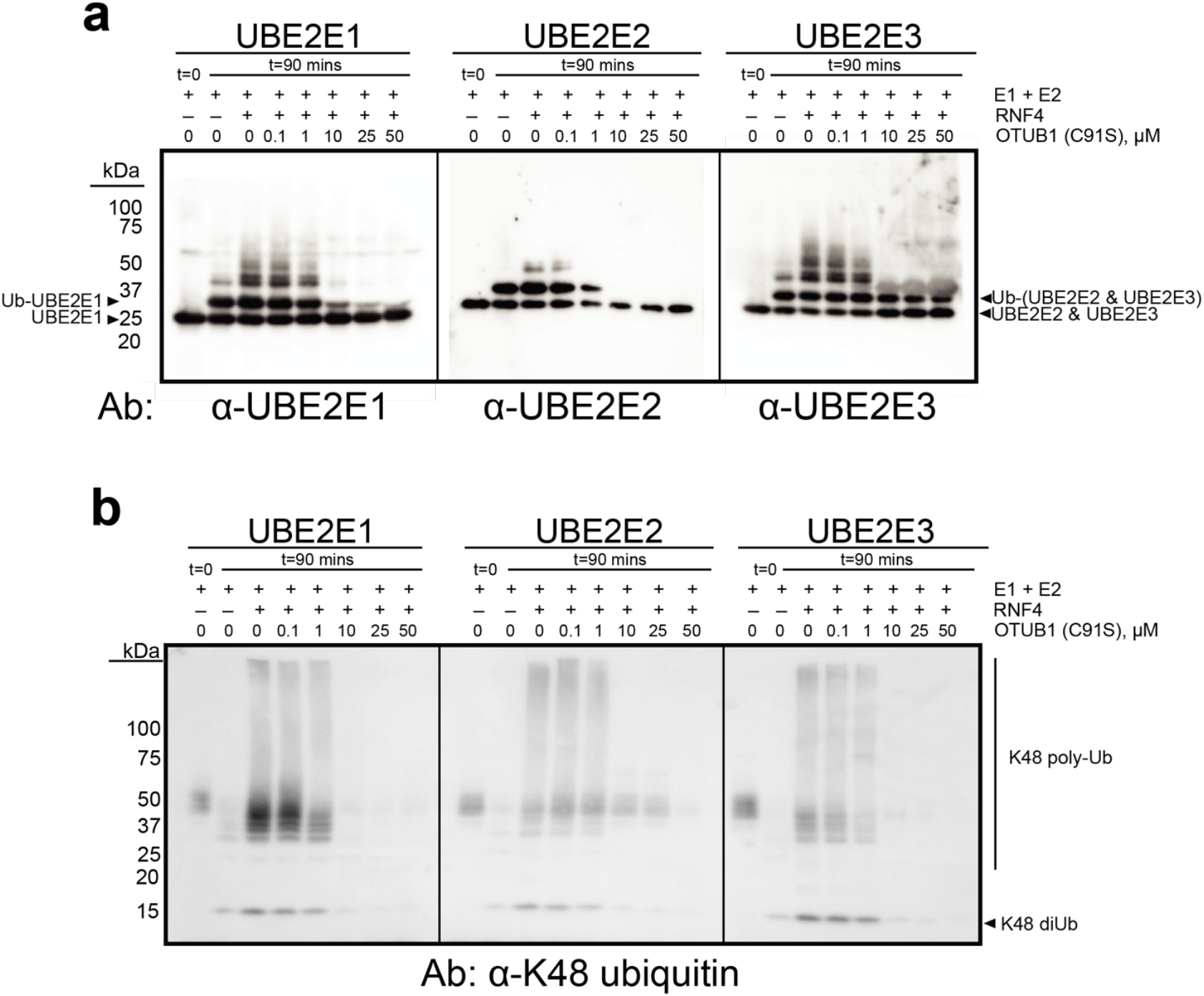
OTUB1 inhibits autoubiquitylation all three UBE2E isoforms. (a) End point reactions containing 0.1 μM E1, 2 μM E2, 2 μM RNF4, and 50 μM Ub wt were quenched at t=0 and 90 mins with increasing concentrations of OTUB1 (C91S), 0-50 μM. Immunoblotting against each E2 was done with isoform-specific antibodies. (b) As in (a) with antibody specific for K48 polyubiquitin. Smears around 50 kDa correspond to a cross reacting species. (Coomassie-stained gel shown in Figure S6).

## Discussion

The ability of OTUB1 to bind select E2 enzymes and the divergent consequences of these interactions – inhibition of ubiquitin transfer by the E2 or stimulation of OTUB1 DUB activity – has raised questions about the conditions under which each activity could play a biological role. The total ubiquitin concentration in mammalian cells is estimated to be between 20 - 85 μM depending on the cell type *^38, 47^*. These numbers account for all forms of ubiquitin: free, in polyubiquitin chains, covalently linked via a thioester linkage to E1 or E2 enzymes, and conjugated to proteins as mono- or polyubiquitin. Only 11% of the total amount of ubiquitin accounts for the concentration of all chains found in cells *^38^*, with K48 and K63-linked chains ranking as the more abundant linkage types *^47^*. The K_M_ and K_d_ of OTUB1 alone for K48 diubiquitin, which we measured at 102.5 and 85 μM **(Figures 4a, 5a)**, respectively, are thus much higher than the expected concentration of K48 chains, making it unlikely that OTUB1 functions effectively as a DUB *in vivo* when not bound to an E2.

Our quantitative study of OTUB1 stimulation by its E2 partners provides a basis for further investigating the relative contribution of these E2 enzymes to OTUB1 DUB activity in cells. As was previously shown for UBE2D2 *^37^*, all E2 partners tested lowered the K_M_ of OTUB1 for its substrate **(Figure 4)**, with no effect on k_cat_. The lowering of K_M_ is a direct consequence of E2 binding which increases the affinity of OTUB1 for K48 chains, as we show for UBE2E1, UBE2D3 and UBE2N **(Figure 5b)**. However, the range of higher K_M_ values suggests that OTUB1 complexes with UBE2E1 or UBE2N are less likely to function as DUBs, as their K_M_ for K48 diubiquitin is 24.1 and 64.3 μM, respectively **(Figure 4b)**. These affinities are well above the expected cellular concentrations of K48 polyubiquitin *^38^*. The weak stimulation of OTUB1’s DUB activity by UBE2E2 and UBE2E3 **(Figure 2a)** similarly suggests that these E2s are unlikely to form active DUB complexes with OTUB1 in cells. OTUB1 binding to UBE2E enzymes is therefore more likely to be important for non-catalytic inhibition of these E2 enzymes, as has been reported for UBE2E1 *^35^*. By contrast, the UBE2D isoforms appear more likely to form functional DUB complexes with OTUB1 *in vivo*. These E2 enzymes have EC_50_ values in the single micromolar range and lower the K_M_ of OTUB1 for its substrate to 6.6 – 14 μM. Since the concentration of UBE2D3 alone has been reported to be 1.7 μM in MEF cells *^19^* and the effective concentration of UBE2D3 needed to stimulate the DUB activity of OTUB1 is 1.6 μM **(Figure 3)**, it is likely that OTUB1:UBE2D3 form a stimulated DUB complex in cells. An important variable, though, is the relative proportion of charged, E2~Ub thioester, versus uncharged E2. Only the uncharged E2 stimulates OTUB1 DUB activity at physiological concentrations of ubiquitin *^37^*, whereas the charged E2~Ub drives formation of a repressed complex with OTUB1 *^28, 30, 31^*. Formation of DUB-active OTUB1:E2 complexes therefore depends on the presence of sufficient uncharged E2, which has been observed to vary among cell lines. UBE2D isoforms have been found primarily in the uncharged state in HeLa and U2OS cells *^37^*, which would favor the DUB-active complex. Moreover, it is possible that responses to a variety of stresses cause transient changes in the E2:E2~Ub ratio, which could shift the balance of OTUB1:E2 complexes from a repressed to active state, or vice versa.

The binding and kinetic data presented here can be used to deduce a thermodynamic cycle describing the equilibrium between complexes formed by OTUB1, its E2 partners and the K48-linked diubiquitin substrate **(Figure 7)**. Since the K_M_ values are similar to the K_d_ of OTUB1 for K48 diubiquitin for the E2s assayed **(Figure 4 and 5)**, we use K_M_ as a measure of substrate affinity for all E2s. The affinity (K_d_) of each E2 for OTUB1 bound to substrate can thereby be derived **(Figure 7b)**, resulting in values in the low to sub-millimolar range **(Figure 7c)**. This analysis highlights the potential for K48 polyubiquitin chains to drive association of OTUB1 with its E2 partners at E2 concentrations on the order of 1 μM or less **(Figure 7c)**. Particularly in the case of the UBE2D isoforms, which are present in cells at micromolar concentrations *^18, 19^*, an increase in K48 polyubiquitin could thereby drive formation of OTUB1-E2 complexes that would degrade the chains while raising the amount of available free ubiquitin. Our quantitative analysis provides a basis for further exploring the biological roles of OTUB1-E2 complexes in cells.

**Figure 7.**
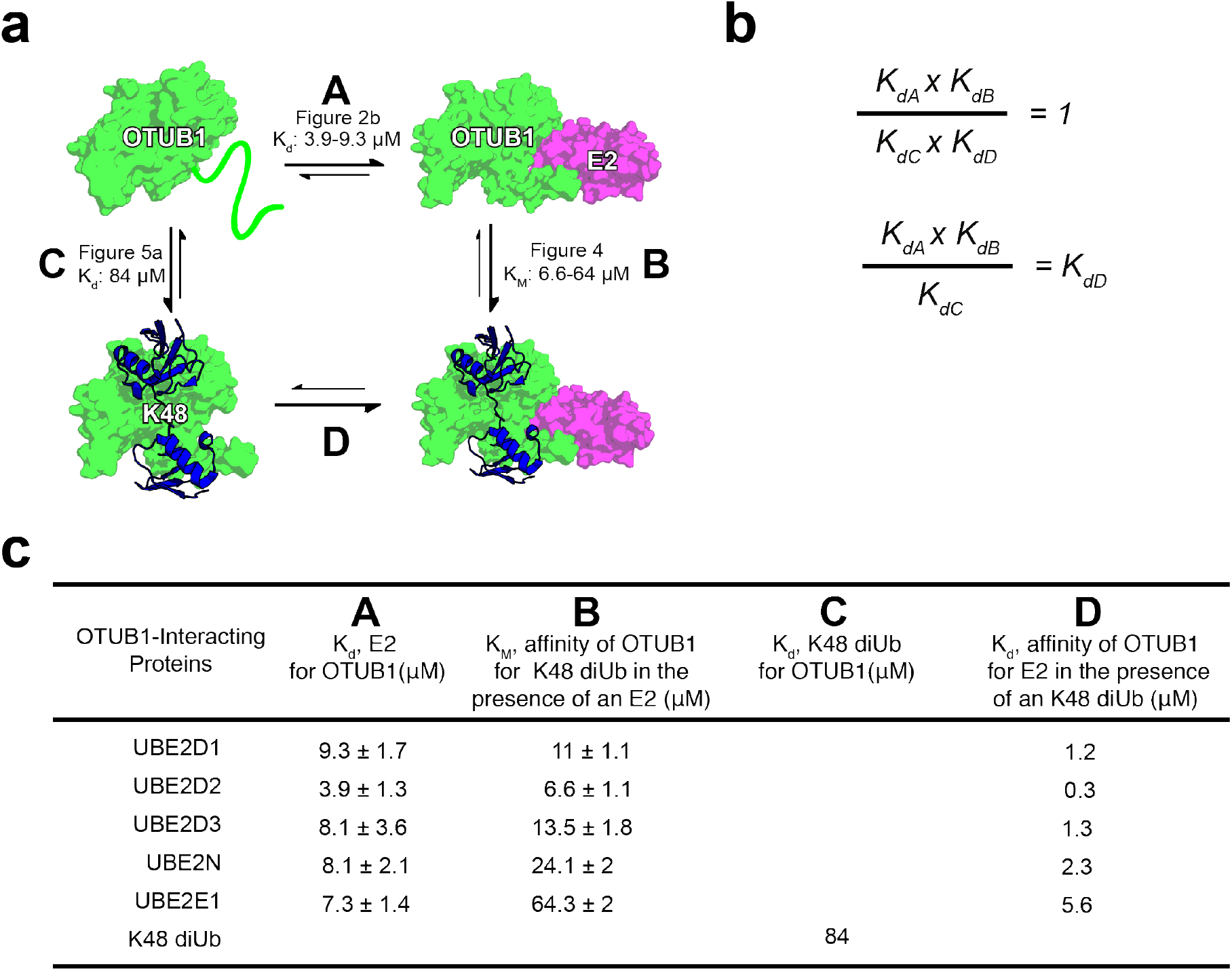
Binding cycle involving OTUB1 interacting with E2 partners and K48 diUbiquitin. (a) Thermodynamic cycle of OTUB1 binding independently to either an E2 or K48 diUbiquitin which in turns favors binding of the other. Each side of the cycle is designated by a letter along with the binding affinities determined in this paper. (b) When comparing each side of the thermodynamic cycle with equilibrium dissociation constants, A x B must equal C x D. Therefore, the constants can be arranged so that K_d_ be calculated. (c) Binding affinities previously determined, A B and C, and those calculated, D.

## SUPPORTING INFORMATION

For additional information, tables and figures, please see Supplemental Info.pdf

## UNIPROT ACESSION CODES

**UNIPROT ACCESSION CODES**
|  |  |
| --- | --- |
| Uba1 | P22314 |
| OTUB1 | Q96FW1 |
| RNF4 | P78317 |
| UBE2D1 | P51668 |
| UBE2D2 | P62837 |
| UBE2D3 | P61077 |
| UBE2N | P61088 |
| UBE2E1 | P51965 |
| UBE2E2 | Q96LR5 |
| UBE2E3 | Q969T4 |
| Cdc34A (UBE2R1) | P49427 |

## ACKNOWLEDGEMENTS

We thank Xiangbin Zhang for synthesizing K48-linked diubiquitin and Evan Worden for critical comments and suggestions. Supported by grants from the National Institute of General Medical Sciences (GM109102 and GM130393).

## AUTHOR CONTRIBUTIONS

L.T.Q. designed experiments, carried out all enzymatic experiments and most ITC studies, performed inhibition studies, and analyzed the data. M.E.M. performed the ITC binding studies for OTUB1 interacting with UBE2E1, UBE2D2, and UBE2N. C.W. supervised the research and assisted with experimental design and interpretation. The manuscript was written through contributions of all authors. All authors have given approval to the final version of the manuscript.

