## Supplemental Figures and Tables for "Characterization of OTUB1 activation and inhibition by different E2 enzymes"

SUPPLEMENTAL INFORMATION

Tables S1 – S3

Figures S1 – S6

| Antibody | Catalogue Number | Supplier | Species | Dilution in 2% BSA, 0.02% Sodium Azide & PBS buffer |
|---|---|---|---|---|
| UBE2E1 | A-630 | Boston Biochem | Rabbit | 1:800 |
| UBE2E2 | Ab-177485 | Abcam | Rabbit | 1:5000 |
| UBE2E3 | PA-551889 | Thermo Fisher Scientific | Rabbit | 1:100 |
| UBE2N | 4E11 | Invitrogen | Mouse | 1:1000 |
| UBE2D | A-615 | Boston Biochem | Rabbit | 1:400 |
| K48 Ub | 4289S | Cell Signaling Technology | Rabbit | 1:1000 |
| Ub wt | sc-8017 | Santa Cruz | Mouse | 1:1000 |

**Table S1. Primary Antibodies.**

| Antibody | Catalogue Number | Supplier | Species | Anti-Species | Dilution in 5% blocking buffer & TBST |
|---|---|---|---|---|---|
| HRP-Conjugate IgG | 12-348 | Millipore | Goat | Rabbit | 1:5000 |
| Alex Fluor 594 conjugate | A-21203 | ThermoScientific | Donkey | Mouse | 1:5000 |

**Table S2. Secondary Antibodies**

| Proteins Interacting with OTUB1 | $K_d$ ($\mu$M) | Stoichiometry (N) | ΔH (kcal/mol) | ΔS (cal/mol) |
|---|---|---|---|---|
| UBE2D1 | 9.3 ± 1.7 | 0.8 ± 0.01 | -3 ± 0.04 | 12.7 |
| UBE2D2 | 3.9 ± 1.3 | 0.9 ± 0.01 | -5 ± 0.12 | 8.5 |
| UBE2D3 | 8.1 ± 3.6 | 1.2 ± 0.03 | -3.8 ± 0.11 | 11.5 |
| UBE2N | 8.1 ± 2.1 | 0.7 ± 0.02 | -5.2 ± 0.16 | 6 |
| UBE2E1 | 7.3 ± 1.4 | 0.9 ± 0.01 | -1.3 ± 0.03 | 19.3 |

**Table S3**. ITC parameters measured by titrating in OTUB1 into E2 binding partners or K48 diUbiquitin

### Figure S1

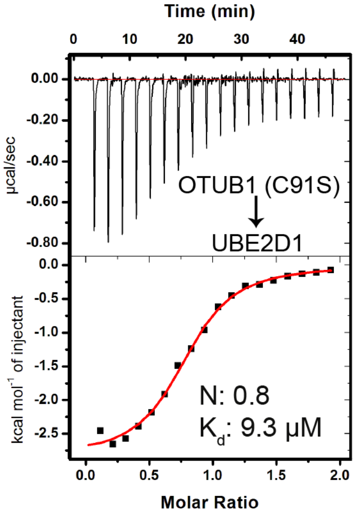


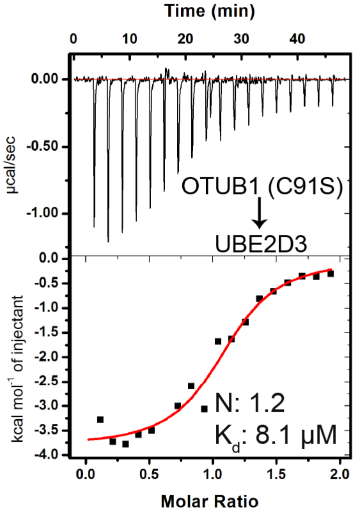


**Figure S1. OTUB1 catalytic mutant binds E2s with the same affinity as wild type OTUB1.** ITC measurement of the affinity of OTUB1-C91S for UBE2D1 (top) and UBE2D3 (bottom).

**Figure S2**

A B

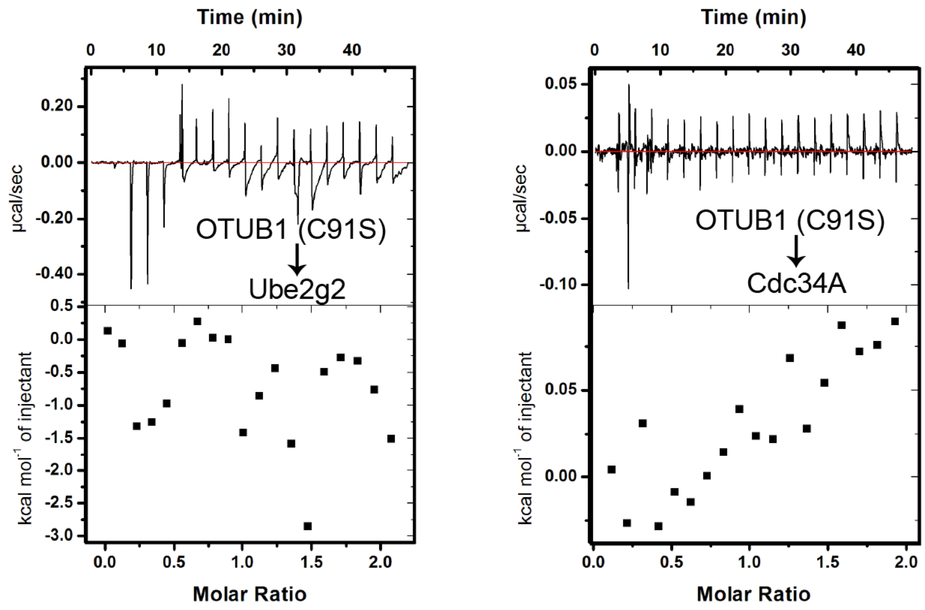


**Figure S2. OTUB1 does not bind to CDC34A or UBE2G2.** ITC measurements in which OTUB1 (1.5 mM) is titrated into a cell containing 150 µM of either Cdc34A or Ube2g2.

**Figure S3**

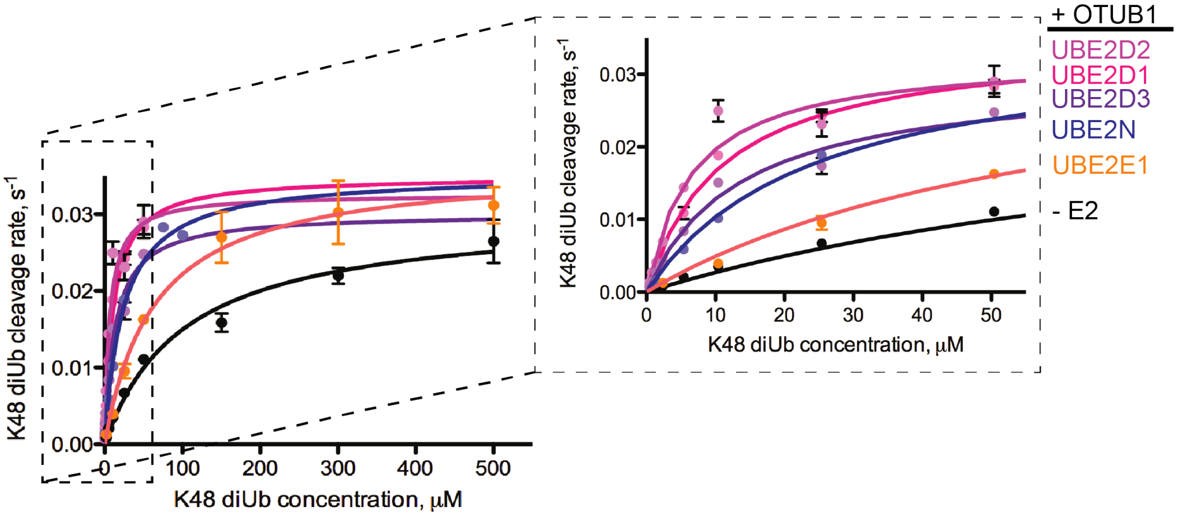


**Figure S3. Full kinetic titration showing effect of E2 enzymes on OTUB1 DUB activity** Full range titration of K48 diUb (0.4 — 500 μM) to track cleavage by OTUB1 (50 nM) in the absence and presence of UBE2D1, UBE2D2, UBE2D3, UBE2N, and UBE2E1 (10 μM). Inset indicates region of plot that is shown in Figure 4a.

Figure S4

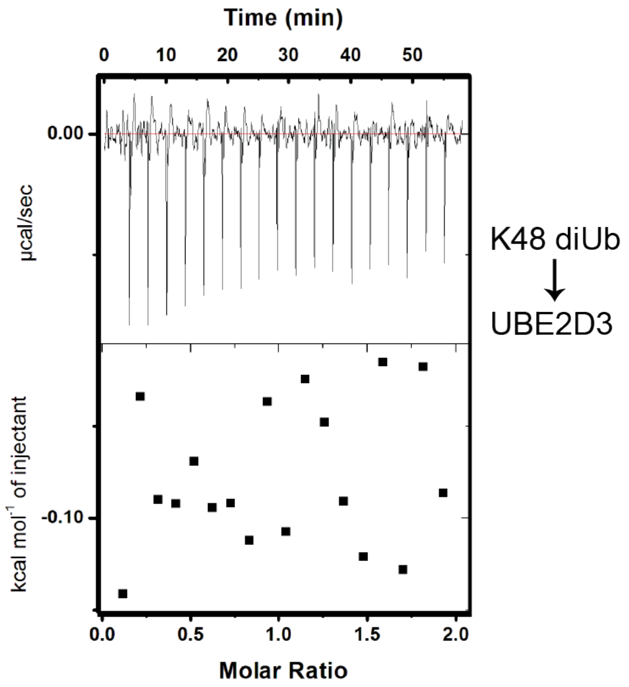


**Figure S4. UBE2E3 does not interact detectably with K48 dibuquitin.** ITC measurements of Binding of UBE2D3 to K48 diubiquitin.

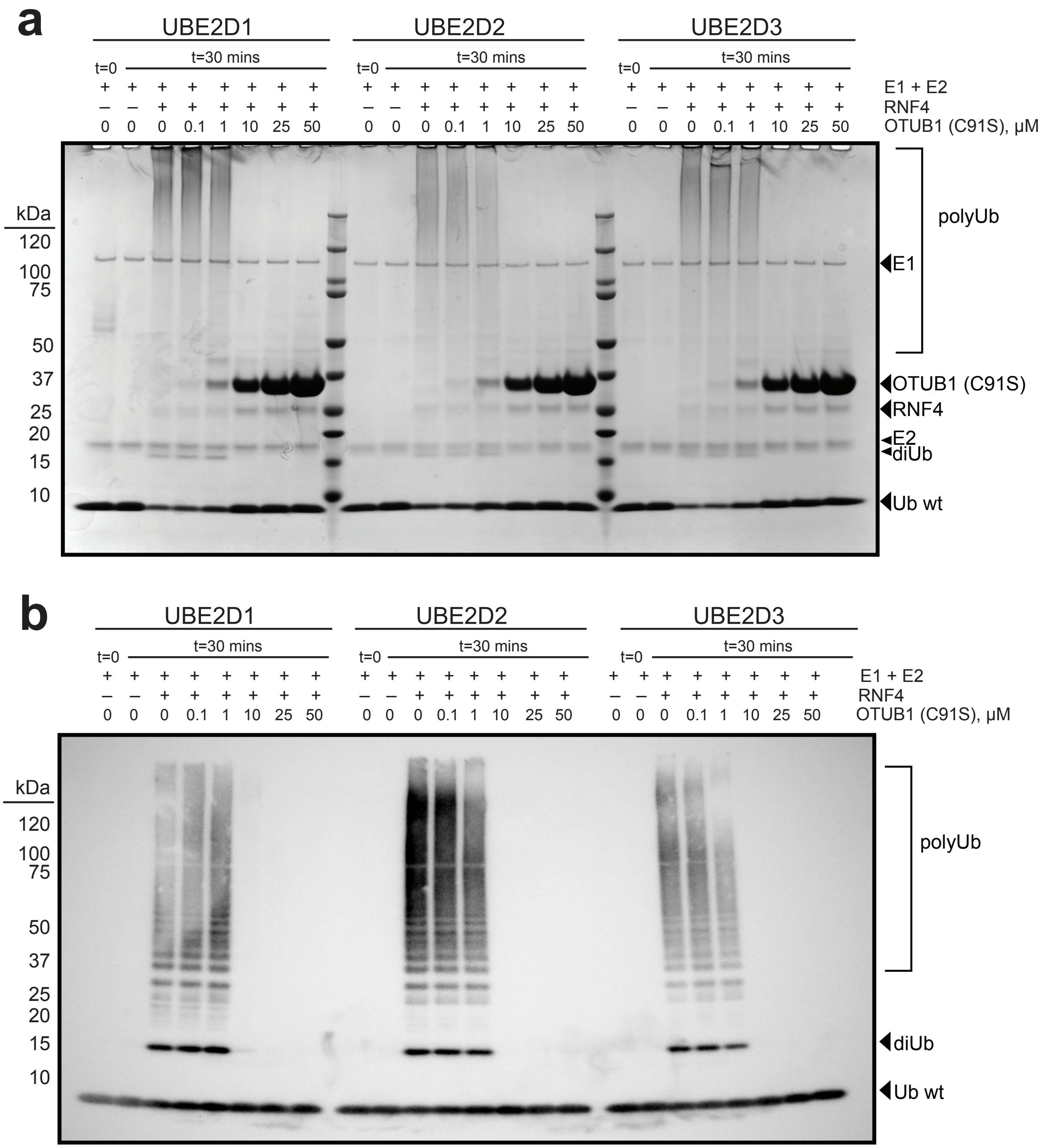


**Figure S5. OTUB1 facilitated inhibition of UBE2D.** End point reactions quenched at t= 0 and 30 mins with increasing concentrations of OTUB1 (C91S) (0-50 μM). Each reaction contained 150 nM E1, 2 μM E2, 50 μM Ubwt, and where applicable, 2 μM E3. **(a)** SDS-PAGE UBE2D(1-3) inhibition by OTUB1 **(b)** SDS-PAGE UBE2D inhibition by OTUB1 transferred to membrane and blotted against Ub wt

### Figure S6

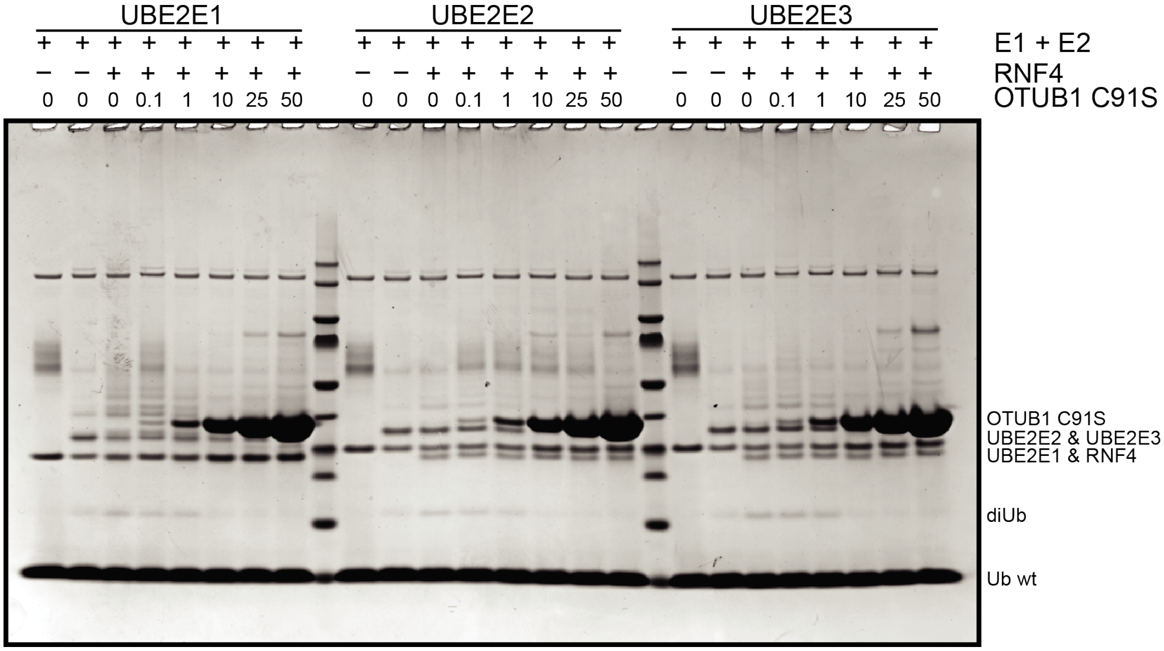


**Figure S6. Coomassie stained gel of OTUB1 facilitated inhibition of UBE2E proteins.** End point reactions quenched at t= 0 and 30 mins with increasing concentrations of OTUB1 (C91S) (0-50 µM). Each reaction contained 150 nM E1, 2 µM E2, 50 µM Ub, and where applicable, 2 µM E3.
